# Network alignment and similarity reveal atlas-based topological differences in structural connectomes

**DOI:** 10.1101/2020.12.16.422501

**Authors:** Matteo Frigo, Emilio Cruciani, David Coudert, Rachid Deriche, Emanuele Natale, Samuel Deslauriers-Gauthier

## Abstract

The interactions between different brain regions can be modeled as a graph, called connectome, whose nodes correspond to parcels from a predefined brain atlas. The edges of the graph encode the strength of the axonal connectivity between regions of the atlas which can be estimated via diffusion Magnetic Resonance Imaging (MRI) tractography. Herein, we aim at providing a novel perspective on the problem of choosing a suitable atlas for structural connectivity studies by assessing how robustly an atlas captures the network topology across different subjects in a homogeneous cohort. We measure this robustness by assessing the alignability of the connectomes, namely the possibility to retrieve graph matchings that provide highly similar graphs. We introduce two novel concepts. First, the graph Jaccard index (GJI), a graph similarity measure based on the well-established Jaccard index between sets; the GJI exhibits natural mathematical properties that are not satisfied by previous approaches. Second, we devise WL-align, a new technique for aligning connectomes obtained by adapting the Weisfeiler-Lehman (WL) graph-isomorphism test. We validated the GJI and WL-align on data from the Human Connectome Project database, inferring a strategy for choosing a suitable parcellation for structural connectivity studies. Code and data are publicly available.

**AUTHOR SUMMARY:** An important part of our current understanding of the structure of the human brain relies on the concept of brain network, which is obtained by looking at how different brain regions are connected with each other. In this paper we present a strategy for choosing a suitable parcellation of the brain for structural connectivity studies by making use of the concepts of network alignment and similarity. To do so, we design a novel similarity measure between weighted networks called graph Jaccard index, and a new network alignment technique called WL-align. By assessing the possibility to retrieve graph matchings that provide highly similar graphs, we show that morphology- and structure-based atlases define brain networks which are more topologically robust across a wide range of resolutions.

## INTRODUCTION

Due to the immense complexity of the brain, it is impossible to gain any insight into its global operation without simplifying assumptions. One such assumption, which has been widely used by neuroscientists, is that the brain, and in particular the cortical surface, can be divided into distinct and homogeneous areas. Of course the definition of homogeneous areas greatly depends on one’s point of view, which has led to a plethora of brain parcellations. For example, the cortical surface has been subdivided based on its cytoarchitecture (Brodmann, 1909), gyri (Desikan et al., 2006), functional organization (Schaefer et al., 2017), axonal connectivity (Gallardo, Wells, Deriche, & Wassermann, 2018), and combinations of these and other features (Glasser et al., 2016). There is also significant evidence that cortical regions vary in shape, size, number, and location across subjects and even across individual tasks, making the existence of a single canonical atlas unlikely. In addition to studying the characteristics of specific brain regions defined by a parcellation, there has been a growing interest in their relationship and interactions, an emerging field known as connectomics. In this context, the focus is shifted from understanding how information is segregated in the brain to how it is integrated. For example, through diffusion Magnetic

Resonance Imaging (MRI) tractography, structural connections between brain areas can be recovered. The result is a network whose nodes correspond to cortical regions and whose edge weights represent the strength of the structural connectivity between pairs of regions. A similar network can also be built from resting state functional MRI yielding a functional, rather than structural, network. These brain networks, which encode the structural and functional connections of the brain, are referred to as connectomes (Hagmann, 2005; Sporns, Tonomi, & Kötter, 2005). Given functional or structural connectomes, their features can be compared across subjects and populations to link network changes to pathology or to further increase our understanding of its organization. An underlying assumption is that a correspondence exists between nodes of the network across subjects, a condition which is usually satisfied by using a group parcellation (Gallardo, Wells, et al., 2018; Parisot, Arslan, Passerat-Palmbach, Wells, & Rueckert, 2015). The drawback of this strategy is that it ignores any subject specific changes in cortical organization and reduces the specificity of the results. An alternative approach is to construct a mapping between the nodes of the network prior to the comparison, therefore allowing the use of subject specific atlases. To our knowledge, this approach has never been investigated in the field of network neuroscience.

The construction of a mapping between network nodes corresponds to what is known in various fields as *network alignment* or *graph matching* (Ayache & Faverjon, 1987; Barak, Chou, Lei, Schramm, & Sheng, 2019; Conte, Foggia, Sansone, & Vento, 2004; Korula & Lattanzi, 2014; Singh, Xu, & Berger, 2008). More recently, the graph matching problem gained attention also in the field of neuroscience, being used in the contexts of the analysis of connectome heterogeneity across subjects (Rasero et al., 2017; Takerkart, Auzias, Thirion, & Ralaivola, 2014) and system-level matching of structural and functional connectomes (Osmanlıoğlu et al., 2019). Graph alignment solutions (called *alignments*) correspond to a permutation of the labels of the nodes of a graph which maximizes its similarity to a second graph. There is no standard way to measure the quality of its solutions (Bayati, Gleich, Saberi, & Wang, 2013). This is also reflected in the neuroimaging literature, where various measures of similarity between brain networks are used (Becker et al., 2018; M. K. Chung, Lee, Solo, Davidson, & Pollak, 2017; Deslauriers-Gauthier, Zucchelli, Frigo, & Deriche, 2020; Osmanlıoğlu et al., 2019; Rasero et al., 2017; Takerkart et al., 2014; Villareal-Haro, Ramirez-Manzanares, & Pichardo-Corpus, 2020). In the context of connectomics, a graph alignment is a reordering of the labels of the nodes of a brain network that maximizes its similarity with a second one while preserving the topology. Describing the brain network through its connectivity (a.k.a. adjacency) matrix, permutations of the node labels correspond to identical permutations of the rows and columns of the connectivity matrix. This problem is distinct from the brain atlas correspondence and parcel matching problems (Gallardo, Gayraud, et al., 2018; Mars et al., 2016). The main difference is that in those problems the permutation acts only on the rows of the connectivity matrix as they find correspondences between *connectivity fingerprints* that rely on external features. Conversely, graph alignment does not rely on any external information and uses only information contained in the topology of the graphs.

The complexity of finding the optimal alignment between two graphs using a naïve brute force strategy is exponential in the number of nodes. It is therefore intractable even for the smallest of brain networks, which typically have 50 cortical regions. Spectral methods are a popular approach to the alignment problem (Feizi et al., 2019; Hayhoe, Barreras, Hassani, & Preciado, 2019; Nassar, Veldt, Mohammadi, Grama, & Gleich, 2018), despite being subject to limitations (Wilson & Zhu, 2008). Modern machine learning paradigms exploit deep learning techniques for finding an alignment (Heimann, Shen, Safavi, & Koutra, 2018; Li et al., 2018; Liu, Cheung, Li, & Liao, 2016), however they make use of partially available information about the alignment itself (Liu et al., 2016), or lack explainability and interpretability.

We first introduce the *graph Jaccard index* (GJI), a natural objective function for the network alignment problem. For a given alignment, the GJI rewards correct matches while simultaneously penalizing mismatches, overcoming limitations of previous approaches (Feizi et al., 2019).

We then propose a new graph alignment heuristic, the *Weisfeiler-Lehman Alignment* (WL-align), based on a weighted variant of the Weisfeiler-Lehman algorithm for graph isomorphism (Weisfeiler & Leman, 1968). WL-align is amenable to concrete interpretability in terms of local network structure around each node (Figure 2) and can be integrated with other heuristics. We compare WL-align against the Fast Approximate Quadratic Programming for Graph Matching (FAQ) (Vogelstein et al., 2015), another efficient brain-alignment heuristic which is solely based on network structure.

## THEORY

A brain network is characterized as an edge-weighted graph *G* = (*V, E*), where each of the *n* nodes represents a brain region and each weight *w*_*ij*_ encodes the strength of the connection between regions *i* and *j*. The graph *G* can always be considered as complete, given that an edge (*i, j*) ∉ *G* can be associated to a null weight *w*_*ij*_ = 0. The matrix that encodes in position (*i, j*) the weight of the edge *w*_*ij*_ between nodes *i* and *j* is called adjacency matrix of *G* and is denoted as Adj(*G*). In the context of connectomics (Hagmann, 2005; Sporns et al., 2005), the adjacency matrix is also known as *connectivity* matrix. In this work we consider only networks with non-negative edge weights. For structural connectomes this does not impose any special preprocessing, since they are usually constructed using streamline count, length, or weights which are already non-negative. However, functional connectomes can contain negative entries because they are typically based on the correlation of resting state functional MRI signals. A practical solution, already used in other studies (Deslauriers-Gauthier et al., 2020), is to threshold the connectomes, therefore replacing negative entries by zeros.

### Brain alignment

To compare two networks, it is of fundamental importance to establish a correspondence between the nodes of the two graphs. Given two networks *G*_1_ = (*V*_1_, *E*_1_) and *G*_2_ = (*V*_2_, *E*_2_) of *n*_1_ and *n*_2_ nodes respectively, it is possible to define an injective map *m* : *V*_1_ → *V*_2_ (whose existence is granted whenever |*V*_1_| ≤ |*V*_2_|) that is called *graph matching* or *network alignment*. An edge (*u, υ*) ∈ *E*_1_ is correctly *matched* by *m* if (*m*(*u*), *m*(*υ*)) ∈ *E*_2_ and both edges have the same weight. Notice that a graph matching that matches all edges corresponds to an injective graph homomorphism. In the context of connectomics we will refer to *m* also as a *brain alignment*. A simple representation of this function is that of a matching matrix *P*_*m*_ of dimension *n*_2_ × *n*_1_ (with *n*_2_ ≥ *n*_1_) defined as

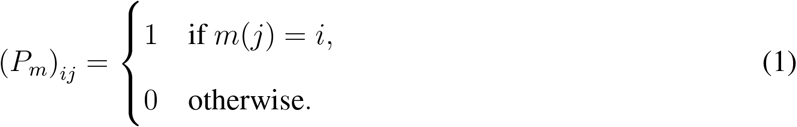

In the special case where *n*_1_ = *n*_2_, *P*_*m*_ is a permutation matrix. If *m* is an isomorphism between *G*_1_ and *G*_2_, then the transformation between the adjacency matrices of the two graphs is fully characterized by the matching matrix and is given by

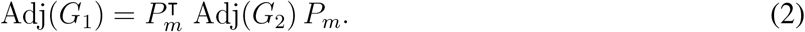

### Quality of brain alignments

Once a brain alignment is identified, its quality can be assessed by evaluating the (dis)similarity of the two resulting networks. On a lexical note, we remark how the concept of *similarity between networks* used throughout this work fits well the standard concept of *matrix similarity* in the particular case where the change of basis matrix is a permutation matrix. In the following, the similarity measures are defined for equal-sized networks, as typically encountered in connectomics. Classical metrics for this task are based on the comparison of the adjacency matrices of the two graphs by means of Pearson’s correlation coefficient, *ℓ*_*p*_ distance, or Frobenius distance (Vogelstein et al., 2015). The norm-based distances estimate the dissimilarity between two graphs *G*_1_ and *G*_2_ by computing the distance between their adjacency matrices as follows

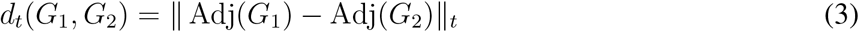

where *t* indicates the type of norm (*p* for *ℓ*^*p*^ norms and *F* for Frobenius norm). Note that higher distance corresponds to lower similarity. Another similarity measure that has been widely adopted in neuroimaging and brain connectivity is correlation; among the many definitions of correlation, we consider

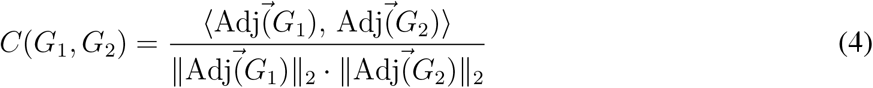

where the numerator is the scalar product between the vectorizations of the adjacency matrices of the two graphs and the denominator is the product of their norms. This similarity measure is also known as *cosine similarity*, since it corresponds to the cosine of the angle between the two vectors. Other distances based on geometrical (Venkatesh, Jaja, & Pessoa, 2020) and homological (M. K. Chung et al., 2017) properties of the networks have been proposed. All such measures capture some aspects of the similarity between two graphs, but none of them satisfies all the following requirements:

- arising as a natural generalization of other similarity measures for less structured data, e.g., for sets of values without a network structure;
- being applicable to the algorithmic graph isomorphism and induced subgraph isomorphism problems, as fundamental special cases of the problem of measuring the similarity between two graphs;
- being simple enough so that its value can be easily interpreted;
- giving a straightforward notion of metric in the considered space.

We therefore propose a new measure obtained by generalizing the Jaccard similarity index, a similarity metric widely adopted in data mining, so that algorithmic problems such as induced subgraph isomorphism can be retrieved as special cases. Moreover, while our proposed measure assigns a clear meaning to the correspondence between two edges in two given graphs, it also depends on the global network structure.

#### Weighted graph Jaccard similarity index

The *Jaccard similarity index* was originally proposed in the context of set theory to measure the similarity between two sets *A* and *B*. It is computed as the ratio between the size of their intersection and the size of their union, that is

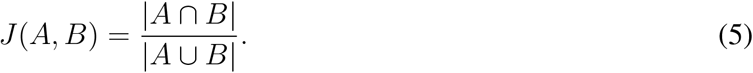

An example of what is measured by the Jaccard index on sets is given in the top panel of Figure 1. Notice that *J*(*A, B*) is defined in the [0, 1] range and the extreme values are attained either when the intersection of the two sets is empty (i.e., *A* ∩ *B* = ∅ ⇒ *J*(*A, B*) = 0) or when the two sets are equal (i.e., *A* = *B* ⇒ *J*(*A, B*) = 1). Both the sets need to be non-empty.

**Figure 1.**
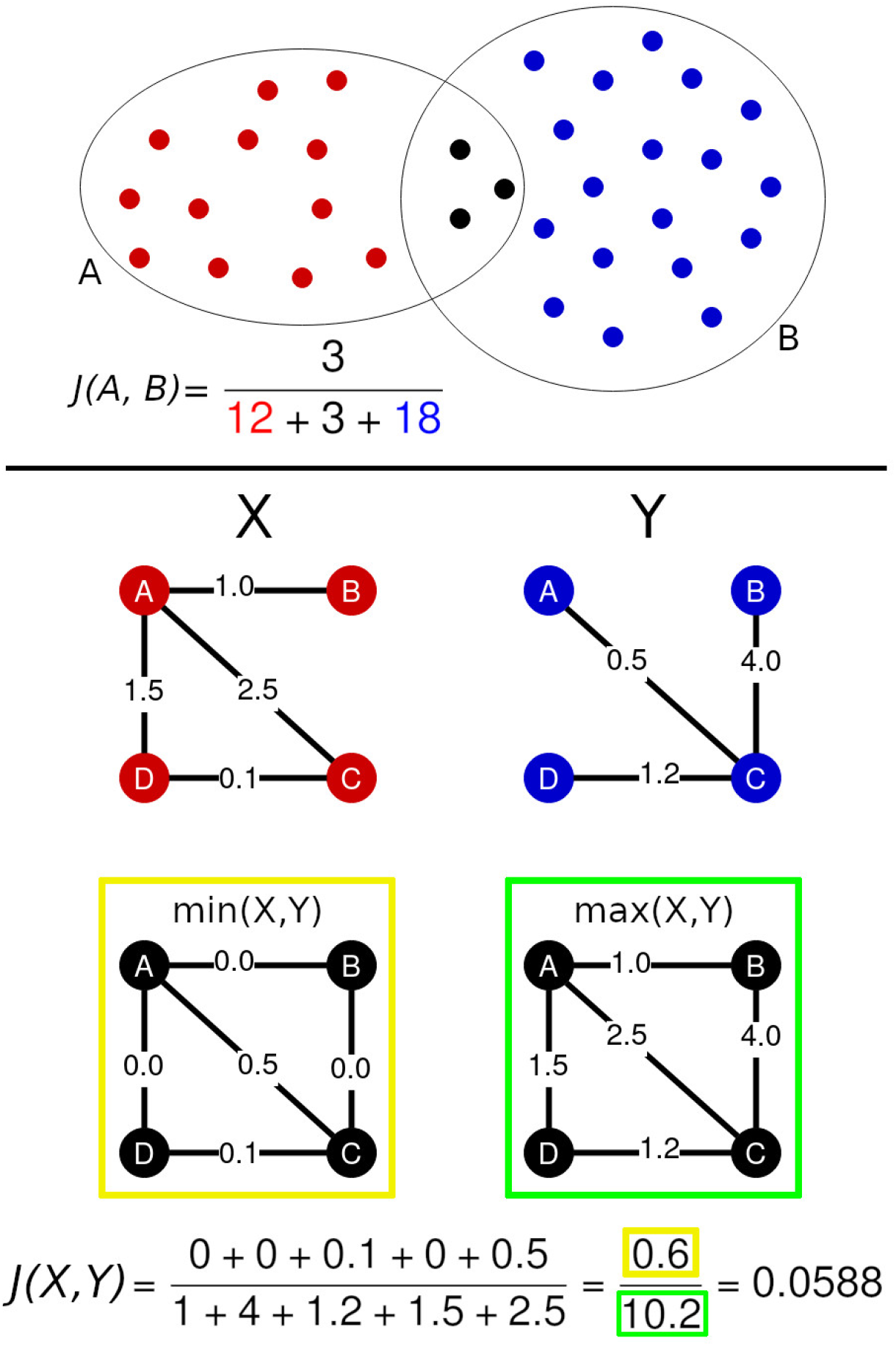
*Top panel.* The two sets contoured by the circles have a non-empty intersection marked by the black dots. The Jaccard similarity index between the two sets is the result of the ratio between the number of elements in the intersection and the number of elements in the union of the two sets. The resulting Jaccard index is equal to *J* = 3/33 ≈ 0.09. *Bottom panel*. This figure shows an example of how to compute the GJI between two compatible graphs *X* and *Y*. For each pair of nodes *i, j* ∈ {*A, B, C, D*}, one computes the minimum and maximum between *X*_*i,j*_ and *Y*_*i,j*_. These two quantities will be used to define the numerator and the denominator of the GJI defined in Equation (7). As shown in the min (yellow) and max (green) graphs, edges that are not in a graph are associated to a null weight. The GJI is then computed as the ratio between the sum of the minimal weights and the sum of the maximal weights.

The Jaccard similarity index has also been generalized to non-negative real vectors and, in this more general setting, is also known as Ruzicka similarity. In detail, given two vectors *x, y* ∈ ℝ^*d*^ such that *x*_*i*_ ≥ 0 and *y*_*i*_ ≥ 0, their *weighted Jaccard similarity index* can be computed as

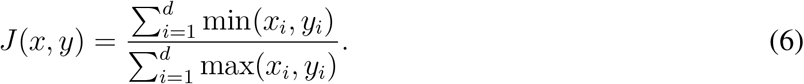

Note that the Jaccard similarity index between two sets follows as a special case whenever the vectors *x, y* are binary and their dimension *d* is equal to the size of the union of the two sets.

Our adaptation of the concept of Jaccard similarity index to weighted graphs is based on the identification of the nodes of the two graphs. Given two brain networks *G*_1_ and *G*_2_ with adjacency matrices Adj(*G*_1_) = *A* and Adj(*G*_2_) = *B*, the *weighted graph Jaccard similarity index* (GJI) of *G*_1_ and *G*_2_ is

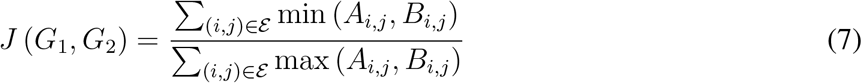

where 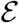 is the set of all possible pairs of nodes. For the sake of the present work, we remark that we can think of *B* as having been previously aligned to *A* via Equation 2. Alternatively, the weighted graph Jaccard similarity index is defined as the weighted Jaccard index of the vectorizations of the graphs’ adjacency matrices. Notice that *J*(*G*_1_, *G*_2_) is not well defined when both *G*_1_ and *G*_2_ are empty (i.e., *E*_1_ = *E*_2_ = ∅). Whenever Adj(*G*_1_) = Adj(*G*_2_), the min and the max in Equation (7) coincide and *J*(*G*_1_, *G*_2_) = 1. On the contrary, if *G*_1_ and *G*_2_ do not have any edge in common (i.e., *E*_1_ ∩ *E*_2_ = ∅), the numerator of Equation (7) will be equal to zero and *J*(*G*_1_, *G*_2_) = 0. A remarkable property of the weighted Jaccard similarity index is that it induces a metric in the space where it is defined. As a matter of fact, the function *d*_*J*_(*x, y*) = 1 − *J*(*x, y*) ∈ [0, 1] respects the three properties of metrics:

1. Identity: *d*_*J*_(*x, y*) = 0 if and only if *x* = *y*;
2. Symmetry: *d*_*J*_(*x, y*) = *d*_*J*_(*y, x*);
3. Triangle inequality: *d*_*J*_(*x, y*) ≤ *d*_*J*_(*x, z*) + *d*_*J*_(*z, y*).

The first two properties trivially follow from the definition of *J*, while the third follows as a particular case of what is presented in (Charikar, 2002, Lemma 1). An example of how the GJI acts on two graphs is given in the bottom panel of Figure 1.

We have so far formally established the notion of network alignment (Equation (1)), and the presented graph Jaccard index as a principled way to measure the quality of an alignment (Equation (7)). We are thus ready, in the next section, to describe our variant of the Weisfeiler-Lehman heuristic and to show how to employ it to construct a network alignment.

### Weisfeiler-Lehman network alignment

In this work we propose a brain alignment technique that allows to define the graph matching *m* between two brain networks *G*_1_ and *G*_2_ with a three-step procedure:

1. For each node *u* in both graphs, define a vector *H*_*u*_ called *signature*.
2. Define a *complete bipartite graph* where on one side there are the nodes of the first graph and on the other side there are the nodes of the second graph; the euclidean distance between two signatures becomes the weight of each edge of the bipartite graph.
3. The graph matching is given by the solution of the *minimum weight bipartite matching problem*, also known as *assignment problem*, on the bipartite graph previously defined.

The novelty element of this brain alignment algorithm is given by the definition of the node signature, which is determined with an algorithm inspired by the Weisfeiler-Lehman (WL) method for graph isomorphism testing (Weisfeiler & Leman, 1968). For this reason, *WL-align* is the name we propose for our brain alignment algorithm.

#### Node signature

The signature that we associate to each node of the two graphs describes the local connectivity pattern of the node. It relies on the concept of *volume* of a node, which is defined as the sum of the weights of the edges incident to the node itself, namely

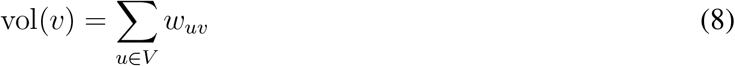

where *υ* is the set of nodes in the graph, *υ* is the node of which we compute the volume vol(*υ*) and *w*_*uυ*_ is the weight of the edge connecting nodes *u* and *υ*. The algorithm that defines the signature of node *u* considers the subnetwork *G*′ induced by the nodes that are reachable from *u* in at most *ℓ* hops. At each of these hops, *G*′ retains only the *k* nodes with highest contribution, weighted according to a function of the path that connects them to *u*. In detail, such a contribution is computed via the following function

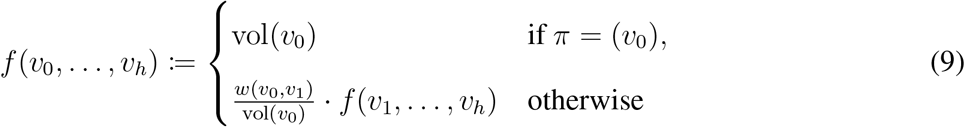

where *w*(*u, υ*) is just a more verbose notation for the edge weight *w*_*uυ*_. The subnetwork *G*′ is a complete *k*-ary tree of depth *ℓ* which can be obtained from a breadth-first search (BFS) starting from *u*, and has a total of 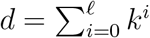 nodes. For this reason the parameters *k* and *ℓ* are respectively called *width* and *depth*. The entries of the signature *H*_*u*_ ∈ ℝ^*d*^ are then computed starting from *u* and following the BFS by recursively, estimating the contribution of each edge to the volume of the considered node via Equation (9). A formal description of the algorithm for computing the signature *H*_*u*_ is given in Algorithm 1, while a graphical intuition is illustrated in Figure 2.

**Algorithm 1.**
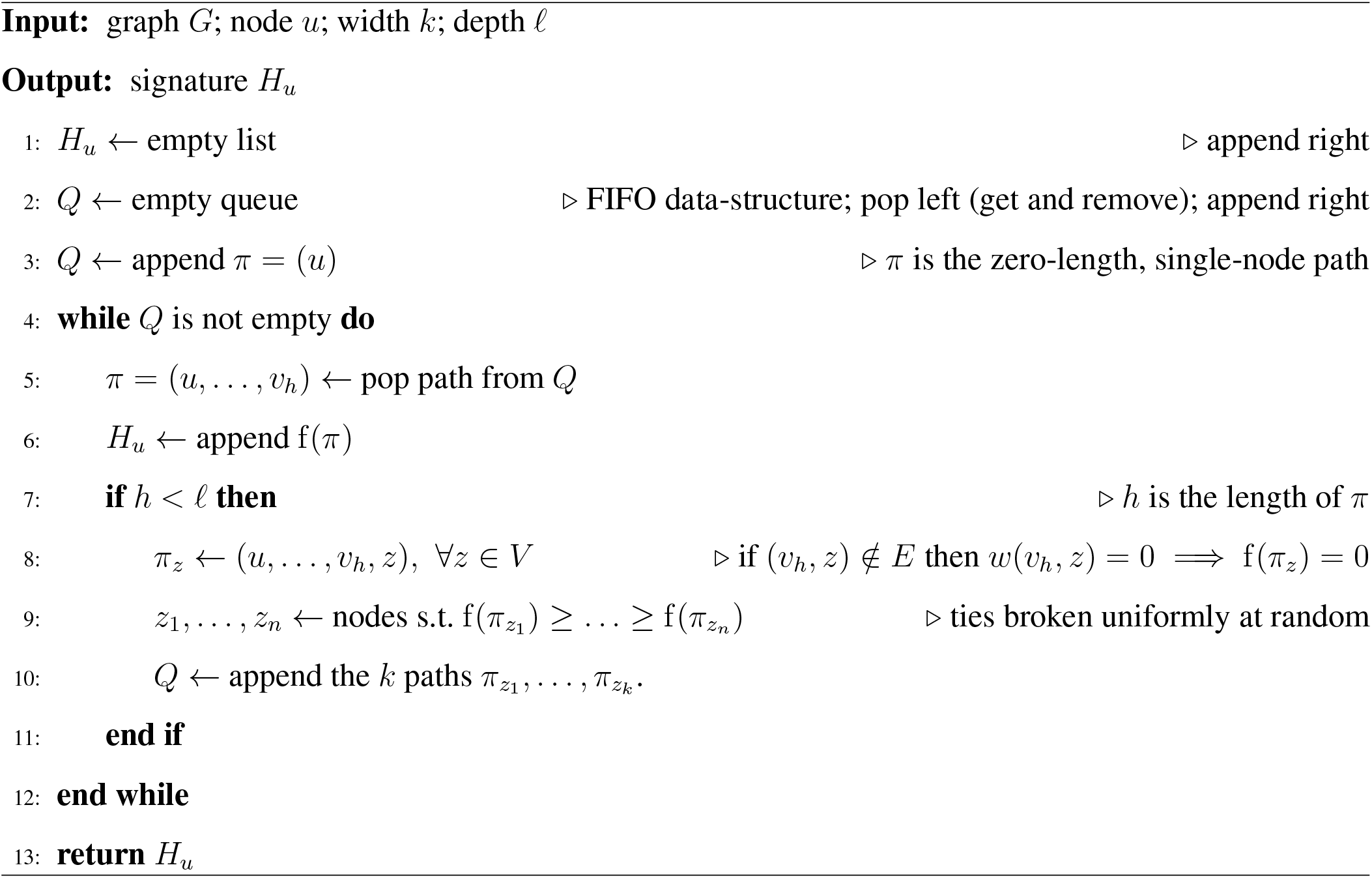
WL-align signature

**Figure 2.**
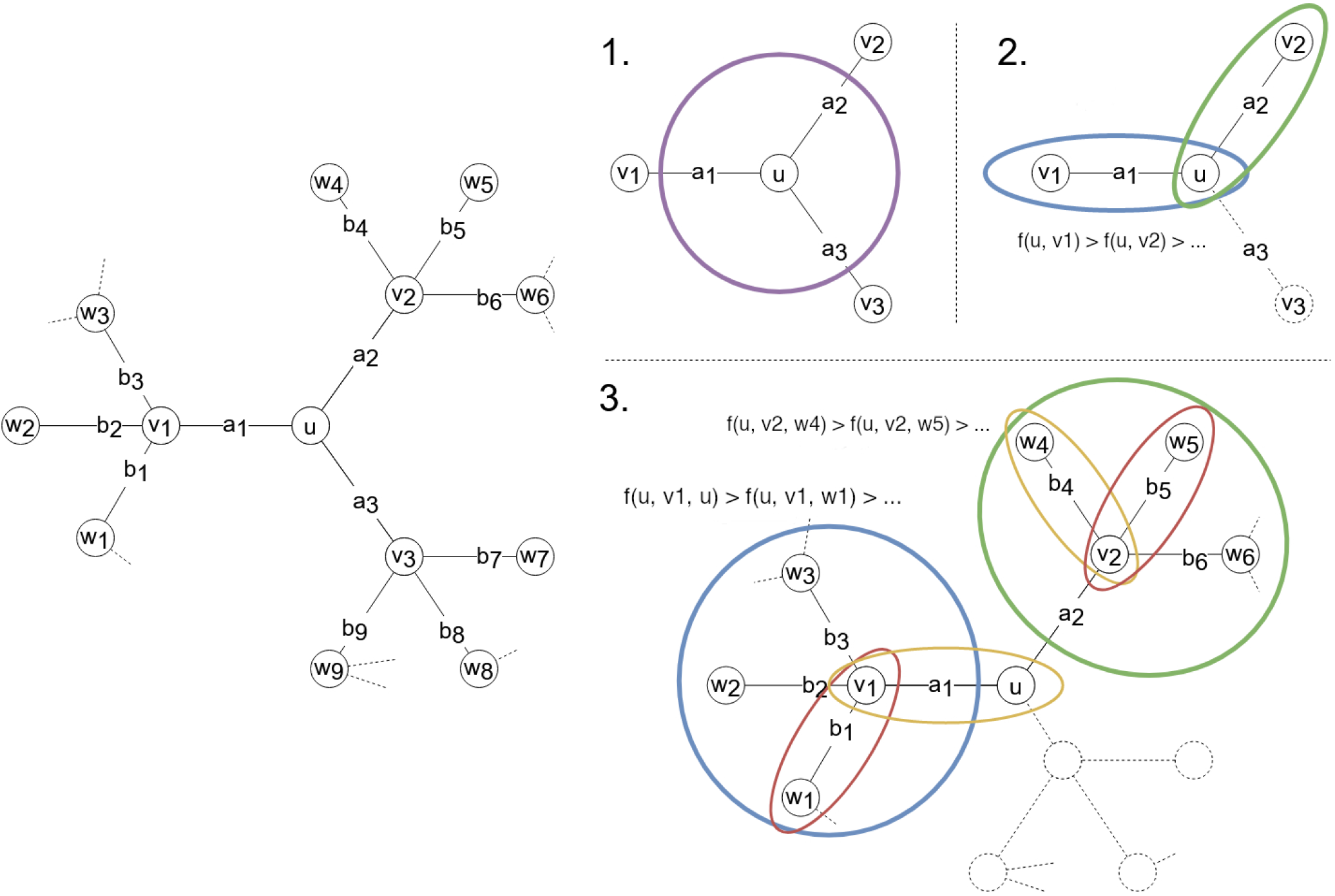
The graph on the left is the one that serves as an example for explaining the algorithm for computing the signature *H*_*u*_ of node *u* with *k* = 2 and *ℓ* = 2. The first entry of *H*_*u*_ is *H*_*u*_[1] = vol(*u*), which is obtained by considering all the edges touching node *u* contoured by the purple circle in panel 1. The focus moves then to the two neighbors that create a path with highest *f*, namely *υ*_1_ and *υ*_2_, which are marked by the blue and green circles in panel 2. They are considered in decreasing order (w.r.t. the volume) and the corresponding entries are computed with Equation (9). For instance, the second entry of *H*_*u*_ is equal to *H*_*u*_[2] = vol(*υ*_1_) · *a*_1_/*υol*(*u*). The third entry is computed in an analogous way as *H*_*u*_[3] = vol(*υ*_2_) · *a*_2_*/υol*(*u*). This concludes the definition of the first 1 + *k* entries of *H*_*u*_. The following entries are defined by considering first the blue and then the green subnetwork in panel 3. The fourth entry is equal to *H*_*u*_[4] = vol(*u*) · (*a*_1_/ vol(*υ*_1_)) · (*a*_1_/ vol(*u*)) and the fifth is *H*_*u*_[5] = vol(*w*_1_) · (*b*_1_/ vol(*υ*_1_) · (*a*_1_/ vol(*u*)). Analogously, the sixth and the last entry will be *H*_*u*_[6] = vol(*w*_4_) · (*b*_4_/ vol(*υ*_2_)) · (*a*_2_/ vol(*u*)) and *H*_*u*_[7] = vol(*w*_5_) · (*b*_5_/ vol(*υ*_2_)) · (*a*_2_/ vol(*u*)).

#### Bipartite graph

Once a signature is computed for each node of the two graphs, we define a weighted complete bipartite graph *G*_*m*_ = ((*V*_1_ ∪ *V*_2_), (*V*_1_ × *V*_2_)). The nodes on the left, i.e., *V*_1_, represent the nodes of the first graph, while the nodes on the right, i.e., *V*_2_, represent the nodes of the second graph. The edge-weights encode the distance between the signatures of pairs of nodes belonging to different graphs, i.e., each edge (*u, υ*) with *u* ∈ *V*_1_ and *υ* ∈ *V*_2_ is weighted according to a function *b* : *V*_1_ × *V*_2_ → ℝ defined as the euclidean distance between the signatures of the two endpoints *b*(*u, υ*) = ∥*H*_*u*_ − *H*_*υ*_∥_2_. Figure 3 shows a simple example of the defined bipartite graph.

**Figure 3.**
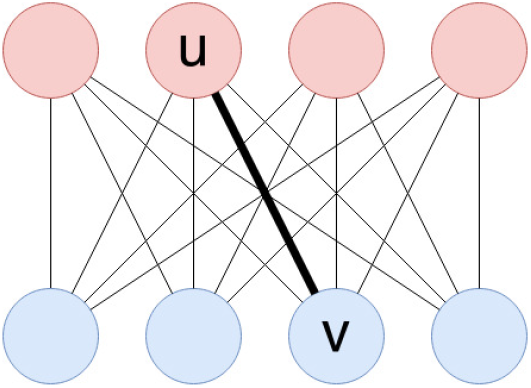
The red and blue nodes in the two rows represent the two graphs *G*_1_ = (*V*_1_, *E*_1_) and *G*_2_ = (*V*_2_, *E*_2_) being aligned. The displayed complete bipartite graph is the one constructed in the second step of the WL-align algorithm. Each edge has weight equal to the euclidean distance between the signatures of the nodes that it connects. For instance, the weight associated to the edge connecting nodes *u* ∈ *V*_1_ and *υ* ∈ *V*_2_ is ∥*H*_*u*_ − *H*_*υ*_∥_2_, where *H*_*u*_ and *H*_*υ*_ are the signatures of nodes *u* and *υ* defined in the first step of the WL-align algorithm.

#### Assignment problem

The final step towards finding the wanted matching with WL-align is the resolution of the assignment problem corresponding to the bipartite graph *G*_*m*_ defined in the previous paragraph. The matching can be found by selecting a minimum-weight graph matching, namely a subset of edges of the bipartite graph such that every node has degree 1 and the sum of the weights of all edges of the subset is minimal. In formal terms, given the two sets *V*_1_ and *V*_2_ and the weighting function *b* that define *G*_*m*_, the problem asks to find a bijection *m* : *V*_1_ → *V*_2_, i.e., the matching, that minimizes the function 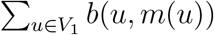. This assignment problem is efficiently solved by the Hungarian algorithm (Jacobi, 1890; Kuhn, 1955).

## METHODS

We processed the data of 100 unrelated subjects from the Human Connectome Project (HCP) database and obtained the structural brain networks via dMRI-based tractography. For each of the 100 subjects we considered 23 parcellations (Desikan, Glasser, Gallardo at 11 different resolutions, Schaefer at 10 different resolutions), obtaining a total of 2300 weighted graphs. For each parcellation, we retrieved a network alignment between each of the 5050 pairs of subjects using WL-align, which is the novel technique introduced in this work, and the state-of-the-art competitor FAQ for a total of 232300 alignments. The quality of the obtained alignments was then assessed using four network similarity measures.

### Data and preprocessing

To build the structural brain networks, we considered the preprocessed data of the HCP database (U100 subject group) (Glasser et al., 2013; Van Essen et al., 2012; WU-Minn Human Connectome Project consortium, 2017). For each subject, a five-tissue-type image (Smith, Tournier, Calamante, & Connelly, 2012) was obtained using the Freesurfer pipeline (Fischl, 2012) invoked through Mrtrix3 (Tournier et al., 2019). A response function was estimated for the white matter, gray matter, and cerebrospinal fluid using a maximal spherial harmonic order of 8 for all tissues (Jeurissen, Tournier, Dhollander, Connelly, & Sijbers, 2014). The fiber orientation distribution functions (fODFs) were then computed using the multi-shell multi-tissue constrained spherical deconvolution algorithm (Jeurissen et al., 2014). Finally, the fODFs were used as input for probabilistic anatomically constrained tractography performed with the iFOD2 algorithm (Smith et al., 2012) seeding from the gray matter - white matter interface and obtaining a total of five million streamlines per subject.

#### Parcellations

The four parcellations considered in this work subdivide the cerebral cortex following different characteristics of the brain. The Desikan (Desikan et al., 2006) parcellation is based on the manual segmentation of a template of the brain cortex that takes into account the morphological consistencies of healthy human brains. For each subject, it was obtained directly from the HCP database (*aparc+aseg.nii.gz*) together with the cortical surface in fslr32k space. The Glasser parcellation (Glasser et al., 2016) follows a multi-modal approach that considers cortical architecture, function, connectivity, and topography. Its projection onto the fslr32k space was obtained from the BALSA repository (Van Essen et al., 2017). The Gallardo parcellation (Gallardo, Wells, et al., 2018) is based on the segmentation of the structural connectivity profiles associated to each point of the cortical surface and the Schaefer parcellation (Schaefer et al., 2017) is based on the analysis of the co-activation patterns of the brain by means of the analysis of resting-state functional connectivity. The Gallardo and the Schaefer parcellations were computed with a granularity of 100, 200, 300, 400, 500, 600, 700, 800, 900, and 1000 parcels. The Gallardo parcellation was computed also with a granularity of 50 parcels. We extracted the 11 Gallardo atlases from the extrinsic connectivity parcellation of Gallardo et al. (Gallardo, Wells, et al., 2018). The used Schaefer atlas (Schaefer et al., 2017) was downloaded from the repository of the CBIG laboratory (Yeo, 2020) for the *seven-networks* parcellation (Yeo et al., 2011). The use of multi-resolution parcellations reflects the multi-scale nature of the brain network and allows to inspect how the atlas resolution affects the similarity and the alignment of brain networks.

#### Connectomes

For each subject and parcellation an in-house software was used for counting the number of streamlines connecting each pair of regions. The obtained quantity was encoded as the weight of the edge connecting the two parcels in the brain network. All the edge weights were then divided by the sum of all the weights in the graph. A total of 23 connectomes of different sizes was obtained for each subject. Given the limitations of dMRI-based tractography, self-connections were excluded from the connectomes, i.e., the diagonal of the adjacency matrix is set to zero. Because of the high resolution of some parcellations, some regions turned out to be isolated (i.e., not connected to any other region). In order to have a connected graph, which is a requirement of the WL-align algorithm, we artificially connected these isolated (i.e., zero-volume) nodes to the others by adding small-weighted edges connecting each of these nodes to all the other nodes in the graph. This weight was set to 1 (before normalization), which from the point of view of tractography is equivalent to the existence of one single streamline connecting the region to the others. The obtained graphs are undirected and weighted.

### Intra-cohort variability

In order to assess the variability between the brain networks of the subjects in the studied cohort, for each subject we measured the similarity between the connectomes of each pair of subjects with three different similarity metrics: the weighted graph Jaccard index (Equation (7)), the Frobenius norm (Equation (3)) and the correlation (Equation (4)).

### Network alignments

In order to assess the ability of WL-align to retrieve the wanted alignment map, we prepared the dataset in a way that allows to test the quality of the alignment against a known ground truth. In practice, for each parcellation *p*, we randomly permuted the node labels of the connectomes of all subjects keeping track of the permutation maps. These permutation maps allow to compute the ground truth matching *m*^*^ between each pair of brain networks computed with the same parcellation.

For the same set of brains, we also computed two graph matchings. The first is *m*_*W L*_, which is computed with the proposed WL-align technique. The width and depth parameters of the WL-align algorithm were fixed to *k* = ⌊log_2_ *n*⌋, where *n* is the number of nodes in the considered network (i.e., one hemisphere), and *ℓ* = 2. We limited the width for efficiency reasons (the size of the signature has size bigger than *k*^*ℓ*^, as described in the previous section) and the depth since further increasing it does not lead to substantial gain w.r.t. the quality of the alignments (the deeper the nodes in the search, the smaller the contribution of the nodes to the signature, as described in Equation (9)).

The second is *m*_*F AQ*_, which is computed with the Fast Approximate Quadratic Programming for Graph Matching (FAQ) algorithm (Vogelstein et al., 2015), which is the state-of-the-art technique for network alignment. FAQ works in three main steps: i) arbitrarily choose a starting bi-stochastic matrix, which acts as a relaxed permutation matrix that aligns the two networks; ii) find a local solution to the Relaxed Quadratic Assignment Problem (rQAP), a dual version of the graph matching problem; iii) project back onto the set of permutation matrices. The solution found by FAQ transforms the adjacency matrix of the first graph into one with approximately minimal Frobenius distance from the adjacency matrix of the second graph. Notice that optimality with respect to the Frobenius distance might not correspond to absolute optimality. We used the implementation of FAQ available in the *graspologic* package (J. Chung et al., 2019) (https://graspologic.readthedocs.io/), setting the number of random initializations to 30.

Both WL-align and FAQ were run separately on each hemisphere of the brain and the two resulting partial alignments were then combined into a single one. The motivation for this choice is that the correct hemisphere can always be assigned to a cortical region, and this property is independent from any influence potentially caused by the registration of the template atlas onto the subject-specific cortical mesh, while other properties, e.g., the location of a region, would be. Moreover, by studying single-hemisphere alignments we bypass the issue concerning the high degree of left-right similarity that characterizes the brain, which could drive the solution towards sub-optimal alignments that are hardly distinguishable without external criteria such as the localization or geometry of the brain regions. Notice that this choice concerns the design of the experiment, not the setup of the graph matching algorithm, which could still be obtained using the full brain network, hence including the inter-hemispheric connections.

### Quality of alignments

Given two networks *G*_1_ = (*V*_1_, *E*_1_) and *G*_2_ = (*V*_2_, *E*_2_) defined on the same parcellation and given a matching *m* between them, we consider the following metrics to evaluate the quality of the matching *m*.

- Node Matching ratio (NMr): the fraction of nodes that have been correctly matched by *m* with respect to the ground truth matching *m*^*^ (known a priori), namely

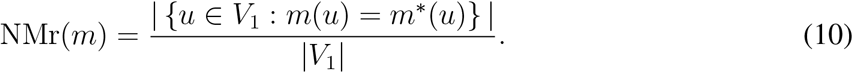 The NMr metric is defined in the [0, 1] range and higher values correspond to better alignments.
- Graph Jaccard index *J* : as defined in Equation (7), namely

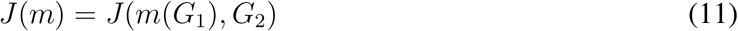

where, with an abuse of notation, we write *m*(*G*_1_) to denote the relabeling of the nodes obtained by applying the matching *m* on the nodes of *G*_1_. Recall that the graph Jaccard index is defined in the [0, 1] range and higher values correspond to better alignment.
- *J*-ratio (*Jr*): the ratio between the graph Jaccard index *J*(*m*) obtained by *m* and the graph Jaccard index *J*(*m*^*^) obtained by the ground truth matching *m*^*^, namely

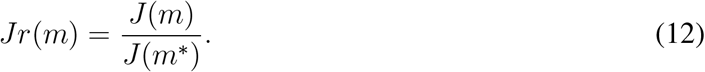 When the ground truth matching *m*^*^ is also an optimal matching, the denominator *J*(*m*^*^) acts as a normalization factor, which takes into account how complex it is to retrieve the matching *m*^*^ in terms of Jaccard similarity; under such assumption of ground-truth optimality, the Jr metric takes value in the [0, 1] range, where higher values correspond to better alignment.
- Frobenius norm (FRO): the Frobenius norm of the difference between the adjacency matrices of *m*(*G*_1_) and *G*_2_, namely

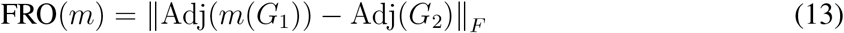

where, as also done for *J*, we write *m*(*G*_1_) to denote the relabeling of the nodes obtained by applying the matching *m* on the nodes of *G*_1_. The FRO metric is defined in the [0, 2] range (since the adjacency matrices both have norm 1) and lower values correspond to better alignment.

For each considered parcellation *p* and for each network alignment algorithm of interest *x* (either WL-align or FAQ), we report the average similarity metric, computed among all pairs of brains in the parcellation. For example, considering NMr as similarity metric, we compute

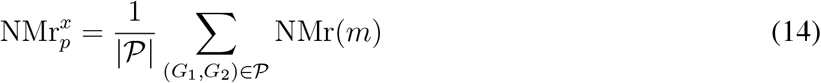

where 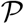 is the set of all pairs of brains with parcellation *p* and *m* is the matching found by algorithm *x* for the input pair of graphs *G*_1_, *G*_2_. Analogously, this is done for all similarity metrics.

A further qualitative assessment of the accuracy of the alignments obtained with WL-align was performed by projecting the matching ratio of each node onto the cortical surface of a randomly picked subject, obtaining a visual indication of the localization of the regions that have been more or less frequently correctly matched. Projecting this information directly on the cortical surface provides insights into the spatial organization of the errors and of the correct matches.

### Statistical analysis

In order to understand the differences between the alignments obtained with WL-align and FAQ, statistical analyses were performed with an alpha of 0.05 in all experiments. A separate analysis was performed for each of the four similarity metrics presented in the previous section. First, for each atlas and pair of subjects we computed an alignment with WL-align and FAQ. For each atlas, we compared the distributions of the values of the similarity metric computed on the alignments obtained with the two techniques using the non-parametric paired-samples Wilcoxon signed-rank test (Wilcoxon, 1945).

## RESULTS

### Experiments

We processed the data of 100 unrelated subjects from the HCP database obtaining the structural brain networks as detailed in the Methods section. For each of the 100 subjects we considered 23 parcellations (Desikan, Glasser, Gallardo x 11, Schaefer x 10), obtaining 2300 weighted graphs. For each parcellation, we retrieved a network alignment between each pair of subjects using WL-align and FAQ. The ability of WL-align to retrieve the correct brain-alignment map was quantitatively evaluated by means of four similarity measures. First, a novel measure of similarity between brain networks called graph Jaccard Index was introduced in the Theory section as an adaptation of the concept of Jaccard index between sets. While behaving in a way which is similar to the commonly used correlation index defined in Equation (4), the graph Jaccard index has the property of defining a metric in the space of connectomes. This is a remarkable property in the context of modern data science, as many standard machine learning techniques can be applied only in metric spaces. The second considered similarity measure is the aforementioned correlation index defined in Equation (4), also known as cosine similarity. The third similarity measure is the Frobenius distance defined in Equation (3), which actually is a dissimilarity measure, therefore connectomes showing higher Frobenius distance are less similar and vice-versa. The node matching ratio defined in Equation (10) is the last considered similarity measure.

### Comparison between similarity measures

Each employed similarity metric answers a specific question. The node matching ratio corresponds to what the expression suggests, namely it counts how many nodes were correctly matched and normalizes the result by the number of nodes in the graph. The other similarity measures have less intuitive definitions. For this reason, and in order to assess the intra-cohort similarity of the connectomes, we measured how much the connectomes of the subjects in the considered datasets are similar to each other with respect to each metric and each parcellation. We recall that the dataset contains only healthy unrelated subjects which do not exhibit any family structure (WU-Minn Human Connectome Project consortium, 2017). This allows to compare how the within-group similarity reacts to the change in resolution and type of the used parcellation. For each parcellation, Figure 4 shows how similar the subjects are with respect to the graph Jaccard index, the Frobenius norm, and correlation. In particular, the figure reports for each parcellation the average similarity across all the pairs of subjects, which can be computed from the ground truth matching that is granted by the fact that each network is defined on a known set of nodes. Despite using the ground truth matching, the graphs are not expected to exhibit perfect similarity (i.e., *J* = 1, FRO = 0 or *C* = 1), as their edge weights are subject-specific. This specificity is what determines the intra-cohort variability that is taken into account into the J ratio similarity metric defined in Equation (12). The most noticeable fact is that the graph Jaccard index and the correlation show an inverted trend with respect to the one of the Frobenius norm. A higher number of parcels gives both lower Jaccard/correlation index and lower Frobenius distance, which a priori is counter-intuitive. This phenomenon is due to the fact that the Frobenius norm is incapable of capturing the relative difference between edge weights and instead considers only the absolute difference between them. As a matter of fact, parcellations with a higher number of parcels will create brain networks with lower edge weights, since the same amount of connectivity (i.e., the same number of streamlines) is distributed among a number of edges that grows quadratically with the number of regions. For this reason, the absolute value of the edge weights will be lower, giving also a lower absolute difference. On the contrary, the graph Jaccard index and the correlation, which are able to capture the relative difference between edge weights, show lower similarity values between brain networks obtained with a higher number of parcels compared to brain networks obtained with a lower number of parcels. This difference suggests that the graph Jaccard index and the correlation mitigate the influence of the number of parcels in the estimation of the similarity between the compared brain networks. Another observation can be done on the singular nature of the Desikan and Glasser parcellations. When measured with the GJI and the correlation, both these parcellations exhibit an intra-cohort similarity in line with the one of the Gallardo parcellation at the corresponding resolutions.

**Figure 4.**
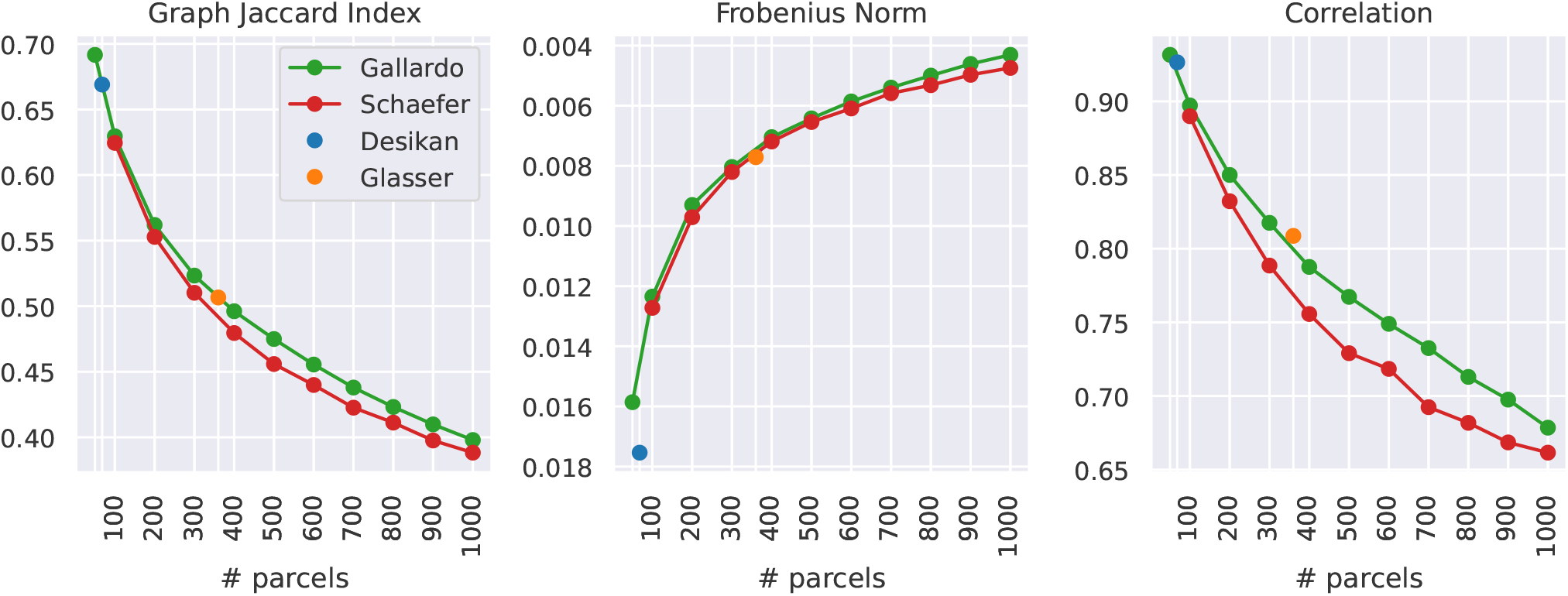
Each point shows the average similarity between every pair of subjects in the considered cohort measured on connectomes obtained with a specific parcellation. The used alignment is the one defined by the ground truth, which in our experiments is known a-priori. All panels show the similarity measure as a function of the number of parcels of the considered atlas. A higher graph Jaccard index and correlation corresponds to higher similarity. On the contrary, a higher Frobenius norm corresponds to lower similarity. In order to keep the intuition that *higher is better*, the *y* axis of the Frobenius norm is flipped.

### Computing brain alignments with WL-align

In this work, the concept of *similarity between networks* was used as a proxy for the quality of a brain alignment, since a good graph matching is expected to correspond to a higher similarity between the aligned graph and the ground truth. A separate analysis was performed for each of the 23 considered parcellations. First, an alignment was computed between each pair of subjects with the proposed technique WL-align and the state-of-the-art algorithm FAQ, then the similarity between the aligned network and the ground truth network was computed with the similarity measures listed in the Methods section. The node matching ratio (NMr) tells the proportion of nodes that were correctly matched by the alignment. This measure does not give any information about the topological differences between the original and the aligned graph, but it gives an important insight on how many nodes are correctly labeled, which may be of fundamental importance in connectomic studies where the regions are associated to a specific function of the brain. The second used metric is the Jaccard similarity index introduced and described in this paper, while the third employed metric is the Jaccard index ratio. The latter measures how the Jaccard index performed with respect to the Jaccard index of the ground-truth matching shown in Figure 4, which is known a priori from the design of the experiment. It differs from the raw Jaccard index in the sense that it takes into account the complexity of the alignment problem, which we showed in the previous section to be more difficult when the number of parcels is higher. A final comparison was made using the Frobenius distance, which is what the FAQ algorithm is designed to minimize. This makes it particularly interesting since we expect FAQ to give Frobenius distance which is less or equal to the one obtained with WL-align.

#### Subject-wise analysis

In the context of this work, the simplest non-trivial alignment to be retrieved is the one between the brain network of a subject and its randomly-permuted version. In this case, a good alignment algorithm is expected to always retrieve the ground truth alignment. In Figure 5 we report the average similarity between the ground truth and the obtained alignment. We notice that WL-align consistently achieves the best possible performance with respect to all the considered metrics. In particular, the naive metric of the node matching ratio always gives similarity equal to 1, meaning that WL-align correctly labels all the nodes whenever a structural brain network is aligned against a randomly-permuted version of itself. These considerations are true for every parcellation. On the contrary, FAQ does not solve the self-alignment problem exactly. All the considered metrics highlight a poor performance of FAQ both in absolute terms and compared to WL-align. As a matter of fact, FAQ on average yields at most 40% of correctly matched nodes, while WL-align consistently gives 100% of correctly matched nodes. Also, different parcellations behave differently when FAQ is employed; for instance, the Desikan parcellation gives lower Frobenius similarity with respect to the other parcellations but shows higher Jaccard index and node matching ratio.

**Figure 5.**
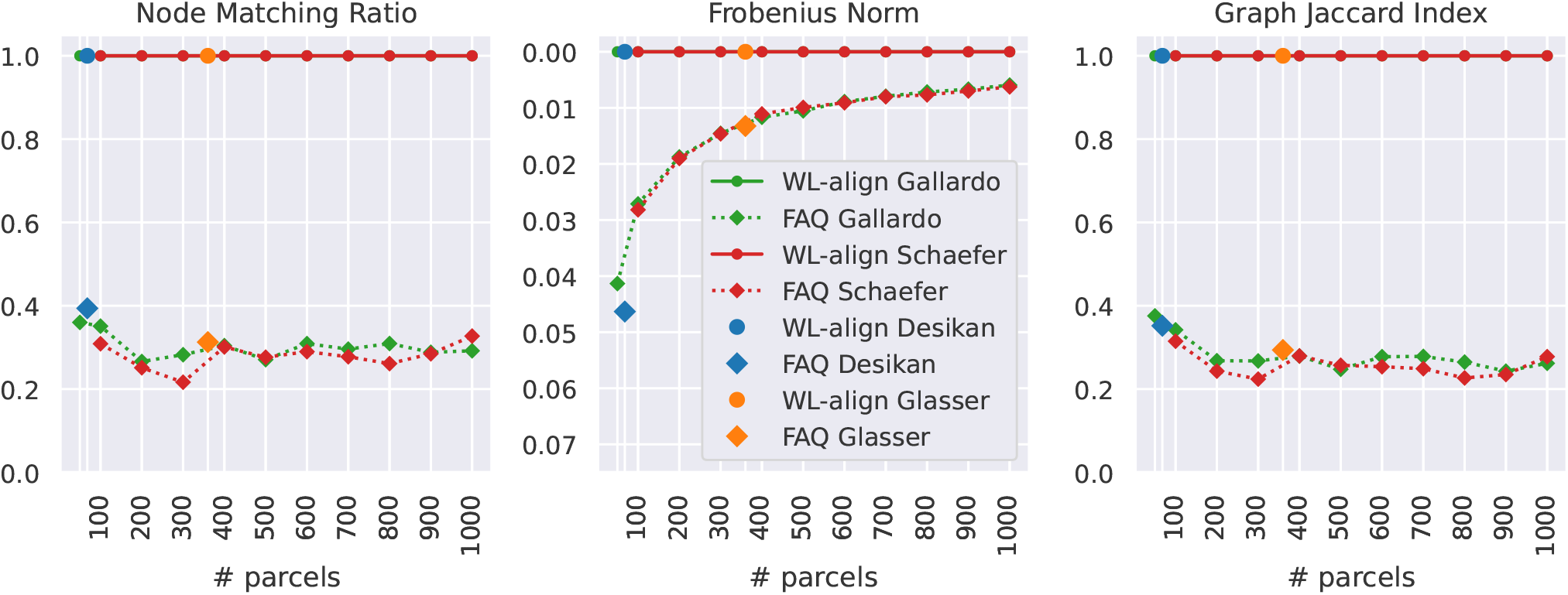
The displayed results concern the alignment between the structural brain network of one subject and its randomly-permuted version. Each panel shows one type of similarity between the aligned networks. Higher values of NMr, Jaccard index and Jaccard index ratio correspond to higher similarity, whereas the Frobenius norm is higher when similarity is lower. In order to keep the intuition that *higher is better*, the *y* axis of the Frobenius norm is flipped. In each panel, one point corresponds to the average (among subjects) similarity computed between brain networks obtained on a specific parcellation and aligned with one technique among WL-align and FAQ. We do not report the results for the *J*-ratio since in this experiment its denominator *J*(*m*^*^) = 1, making the plot identical to the one of the graph Jaccard index. All the four plots show the similarity as a function of the number of parcels in the considered atlas.

#### Full cohort analysis

When all the subject are aligned with the permuted version of each other, the problem is more complicated. Even though we considered healthy subjects whose acquisition followed the same protocol and that have been processed in an identical way, the subject-specific differences and the intrinsic noise of the data yield estimated structural brain networks that are in practice different among each other, despite being substantially coherent. In order to assess the ability of the proposed alignment technique to overcome these differences and yield an alignment as close as possible to the ground truth, we considered all the alignments between each pair of subjects, including the ones between a subject and a randomly-permuted version of itself. The brain alignments obtained with WL-align are compared to the ones computed with FAQ and presented in Figure 6, which reports the average similarity between the obtained alignment and the ground truth alignment among all the possible pairs of subjects. The statistical significance of the differences between results obtained with WL-align and FAQ is assessed using the non-parametric paired-samples Wilcoxon signed-rank test (Wilcoxon, 1945). For the studied cohort, statistically significant differences are observed for each atlas and each employed similarity metric, as shown in Section B of the supplementary material. In terms of Frobenius norm, the alignments obtained with WL-align and FAQ are very similar, with WL-align systematically showing slightly higher Frobenius similarity. The performance of the Gallardo parcellation is indistinguishable from the one of the Schaefer parcellation. Also, the Glasser parcellation is in line with the Schaefer and Gallardo parcellations when the alignment is obtained with WL-align, while this is not true for the Desikan parcellation. Recalling that FAQ is a technique that is inherently based on the Frobenius norm and WL-align is not, we can notice that WL-align gives a brain alignment that does satisfies also the optimality criteria of FAQ, additionally to its own. A second thing that we can notice about the Frobenius norm is that it exhibits the same phenomenon as in Figure 4, where the Frobenius similarity increases with the number of parcels. This phenomenon appears for the same reason as before, namely the Frobenius norm does not capture the relative difference between the edge weights in the compared networks. All the other employed similarity metrics suggest that WL-align has superior performance with respect to FAQ. While FAQ has almost identical performances when applied on the Gallardo and the Schaefer parcellations, WL-align shows relevant and previously unobserved differences between the performances of the two. In particular the Gallardo parcellation allows to retrieve better alignments with respect to the Schaefer parcellation. This may be due to the fact that we are studying structural connectivity, therefore the use of a function-based parcellation like the one of Schaefer may affect the quality of the alignment when compared to the structural connectivity computed on a structure-based parcellation like the one of Gallardo. Looking at the behavior of the Desikan and the Glasser parcellation, we notice two different scenarios. The Glasser parcellation shows Jaccard similarity slightly lower than the one of the Gallardo parcellation but still higher than Schaefer’s, suggesting that the multi-modal nature of the atlas allows to capture, at least in part, the structural connectivity features that we are looking at. This contrast is evident only when WL-align is employed. The Desikan parcellation behaves differently. While exhibiting lower performance with FAQ, when the WL-align is employed it emerges as a slightly superior parcellation with respect to the NMr, the GJI and the J ratio. We finally notice that atlases with *>* 400 parcels all behave very similarly, namely they reach a plateau in terms of Jaccard index, Jaccard index ratio and node matching ratio. This is true both when WL-align and FAQ are employed. The performance in this range is lower than the one in the 50 − 400 parcels range.

**Figure 6.**
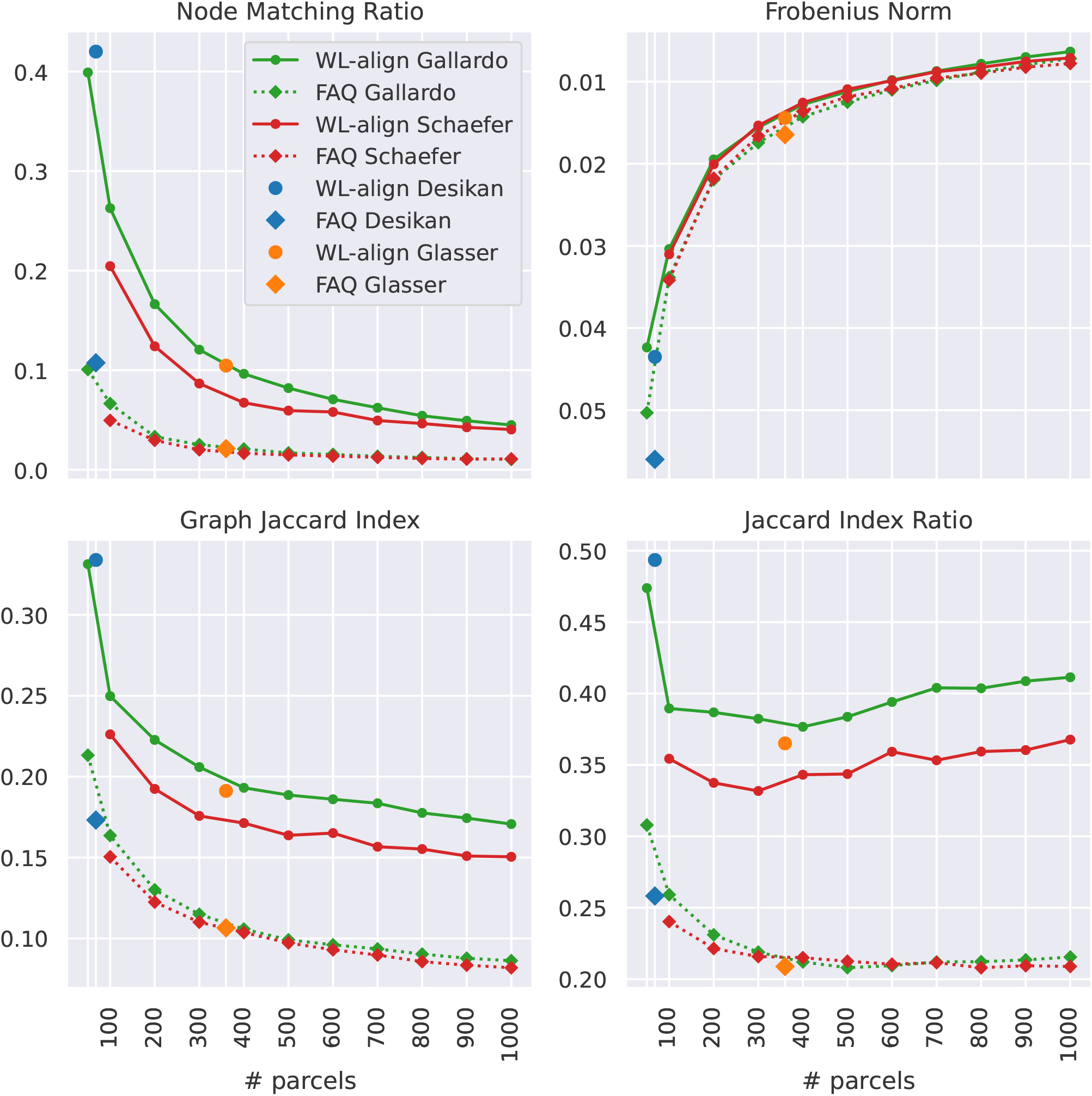
The displayed results concern the alignment between the structural brain networks of each pair of subjects including the self-comparisons. Each panel shows one type of similarity between the aligned brain networks. Higher values of NMr, Jaccard index and Jaccard index ratio correspond to higher similarity, whereas the Frobenius norm is higher when similarity is lower. In order to keep the intuition that *higher is better*, the *y* axis of the Frobenius norm is flipped. In each panel, one point corresponds to the average (among subjects) similarity computed between brain networks obtained on a specific parcellation and aligned with one technique among WL-align and FAQ. All the four plots show the similarity as a function of the number of parcels in the considered atlas.

### Self matching rate

Figure 7 illustrates the self matching rate for each region of 9 example atlases, i.e., the fraction of times regions were correctly matched when aligning different brains represented using the same atlas. It is clear that, as the number of parcels is increased, the matching rate is reduced. This can be explained by the increased difficulty of the alignment problem, but also by a decrease in the signal-to-noise ratio of the connectomes driven by the reduction in parcel size. It is also interesting to note that the matching rate does not appear to be symmetric across hemispheres. For example, the right inferior parietal region of the Desikan atlas obtains relatively high matching rate of roughly 0.8, whereas the contralateral region only obtains roughly 0.4. This analysis gives important insights into the type of errors that are made by WL-align. In particular, it shows that the incorrect matchings do not have a particular structure that can be related to the geometry and morphology of the brain, be it some regional concentration of errors or some consistent symmetry with respect to the hemispheres.

**Figure 7.**
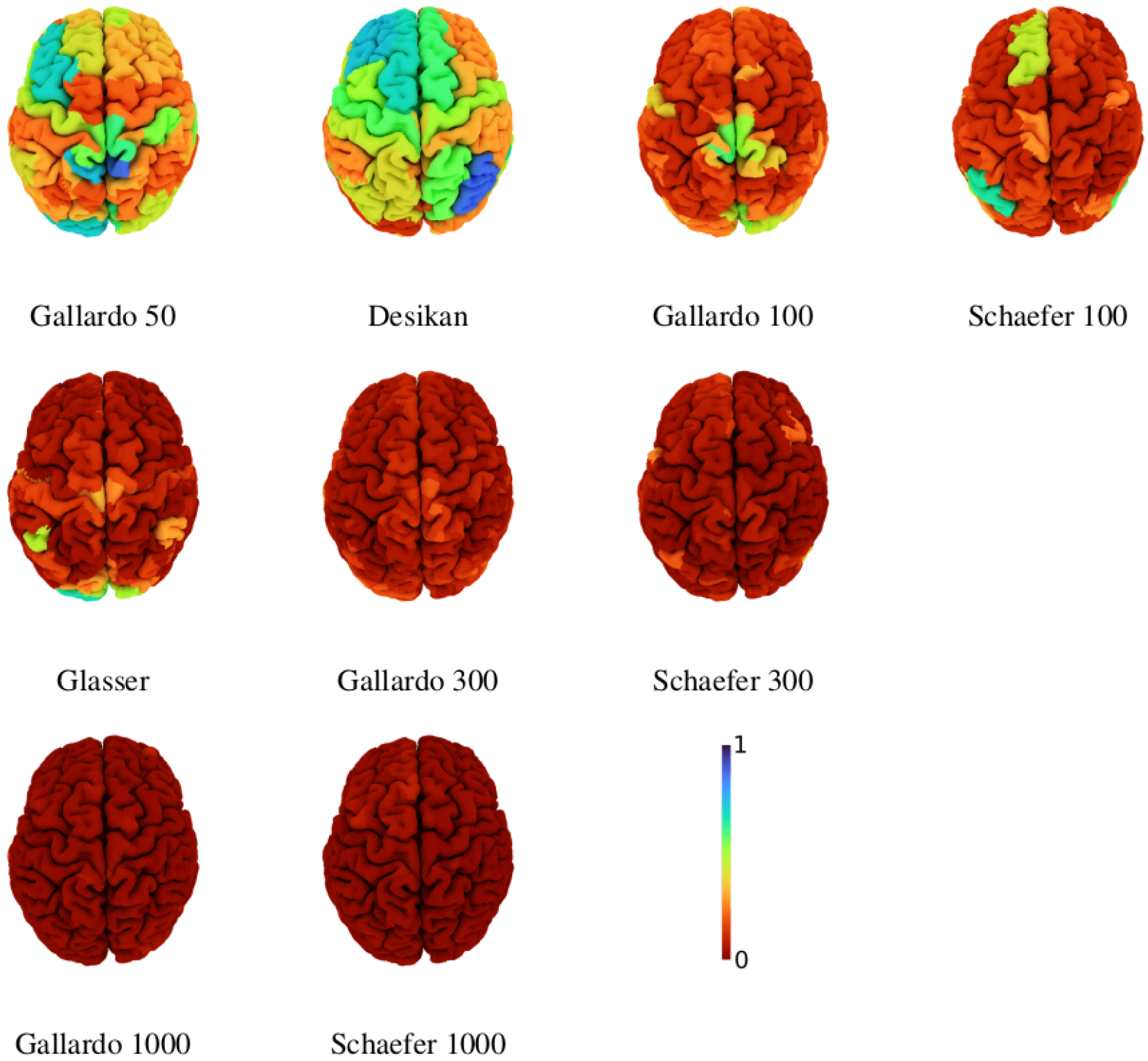
Self matching rate of the labeling per region for different atlases using WL-align. Atlases with 100 regions or less are illustrated in the first row. The second row illustrates atlases with approximately 300 regions and the third row those with 1000 regions.

## DISCUSSION AND CONCLUSIONS

Among the fundamental problems of network neuroscience at the scale of whole-brain structural connectivity, finding correspondences between brain regions and quantitatively assessing the similarity between brain networks are particularly important when it comes to considering massive heterogeneous datasets and modern data science techniques. In this work we considered these two problems in relation with the unresolved question concerning the choice of the parcellation for structural connectivity studies.

We proposed and analyzed a similarity index between brain networks, inspired by the Jaccard index between sets, that behaves in a way similar to the classical correlation index. Additionally, it enjoys the remarkable property of defining a metric in the space of connectomes, which is interesting both from the theoretical point of view and for data science applications. The proposed graph Jaccard index showed to be less affected by the number of regions in the chosen parcellation than the Frobenius distance, which is one example of (dis)similarity index from the class of norm-based distances.

The second object introduced in this paper is WL-align, a novel algorithm that allows to find the graph alignment between two brain networks. It relies solely on topological features of the brain network, which makes it particularly suitable for being applied also outside the domain of network neuroscience. When WL-align is used in our experiments in order to retrieve the alignment between a network and a permuted version of itself, it gives the exact solution. This does not happen when the main competitor FAQ is employed. The superior performance of WL-align is evident also when brain networks of different subjects are aligned. In this case, the WL-align algorithm was shown to retrieve brain alignments that are closer to the ground truth with respect to the alignments obtained by FAQ, and we showed that the difference between the WL-align and the FAQ alignments is statistically significant in the studied population. Notice that as it is designed, the WL-align algorithm builds on the construction of a feature vector for each node of the graph, which is then used as an edge weight in an assignment problem on a bipartite graph. This does not include any prior knowledge other than the topological similarity between the two networks to be aligned. The analysis provided in this work was intentionally constrained to the pure topological comparison of networks. Nevertheless, it would be possible to extend the feature vector defined in WL-align with any prior of geometrical, spatial, anatomical or connectomic nature or to add any constraints in the assignment problem on the bipartite graph. Future works will be devoted to the design of these constraints and features.

The proposed WL-align algorithm can be further adapted to work with types of network other than the structural networks studied in this work, which are undirected and have non-negative edge weights. The most intuitive way to adapt WL-align is to change the way in which the node signatures are defined, then set up the bipartite graph and find the matching with the Hungarian algorithm in the canonical way. A first interesting case is represented by weighted networks having both positive and negative weights. This is the typical case of *functional* connectivity studies, where the connectivity between regions is evaluated as the correlation (i.e., *w*_*ij*_ ∈ [−1, 1]) between the activation in different regions (Van Den Heuvel & Pol, 2010). As we defined it in this work, WL-align would select the most relevant *d* nodes in an unpredictable way due to the presence of negative-valued entries in Equation (9). A possible adaptation of it would be to select the relevant edges performing the breadth-first search ignoring the sign of the weights, then evaluating the corresponding entries of the WL signature using the signed edge weights in Equation (9). Another possible adaptation would require the decomposition of the adjacency matrix of the network as Adj(*A*) = *A*_*p*_ − *A*_*n*_, where *A*_*p*_ is the positive part of the matrix and *A*_*n*_ is the negative part of the matrix. Notice that the graphs corresponding of both *A*_*p*_ and *A*_*n*_ will have non-negative edge weights. For each node, the WL signatures obtained from *A*_*p*_ and *A*_*n*_ can be concatenated, then used in the canonical way. Another interesting case is represented by directed networks, which in the context of brain imaging represent the concept of *effective* connectivity (Friston, 2011). Here, the only further adaptation that would be required is a careful definition of the breadth-first search that gives the selection and the order of nodes that are used for defining the WL signature. For directed positive-weighted network, the algorithm works as it is, while for directed networks with signed weights it would require the adaptations mentioned for the case of functional connectivity. Finally, we discuss the adaptation of WL-align to temporal networks. This type of graph has gained much interest in the context of brain imaging since the concept of *dynamic functional connectome* (Preti, Bolton, & Van De Ville, 2017) has been introduced and the consequent definition of specific tools for the graph-theoretical analysis of these time-dependent networks (Sizemore & Bassett, 2018). In this case, at least two options can be explored: first, one could concatenate the WL signatures of each node obtained at each time point, then run WL-align in the canonical way. Alternatively, it would be possible to perform the breadth-first search by taking into account the temporal component, hence traversing the graph both in space and time.

An important instance of the graph matching problem which we did not consider in this work corresponds to when the two networks that are being aligned have different numbers of nodes. Being a generalization of a graph isomorphism test, WL-align does not appear to be trivially adaptable to this case. A possible solution would be to employ some dimensionality reduction technique (e.g., clustering via community detection) in the larger graph to reduce the number of nodes to the one of the smaller graph, then use WL-align to retrieve the wanted alignment.

Some remarkable conclusions concerning the parcellations to be used in structural brain connectivity studies can be drawn from the ability of WL-align to find the correct alignment between two brain networks. First, the function-based parcellation of Schaefer is a poorer choice than the structure-based parcellation of Gallardo, the multimodal parcellation of Glasser and the morphological parcellation of Desikan. This was expected from the fact that the whole study is centered on measuring *structural* connectivity, hence the choice of a function-based parcellation was never expected to be optimal from any point of view. Allowing to express this concept quantitatively is one of the merits of WL-align. A second remarkable aspect is the performance of the Desikan atlas, which gave better results in terms of alignability than any other parcellation of any granularity. For this reason, whenever a study is designed using a coarse parcellation of the cortex (in the 50-200 parcels range), one should consider using the Desikan atlas as a first choice. Not only it would be a highly reliable choice that has been consistently used throughout time in the community, but with this study we showed that it would also allow to define brain networks with more consistent topological features, in particular those captured by WL-align. As far as brain atlases with a higher number of parcels are concerned, we showed that parcellations with a number of parcels in the > 400 range have lower performance in terms of GJI and NMr. However, when the inter-subject variability is taken into account in the evaluation of the similarity, as for the case of the Jaccard index ratio, we see that the performance is nearly constant for atlases with > 300 parcels.

The change in performance that we observe with the growing resolution of the atlas could also be due to the number of streamlines employed in the tractography pipeline, which could be adapted to the used atlas, but in practice is the same for every atlas at each resolution. On the other hand, the standardized tractography pipeline (including the identical number of streamlines in each tractogram) is what allowed us to present a comprehensive analysis and comparison of the performance across resolutions. In order to disambiguate this point, it would first be necessary to analyze how the strongest connections in the network (hence those considered by WL-align) are affected by the number of tracked streamlines. An alternative solution could be to employ a tractography filtering technique such as SIFT2 (Smith, Tournier, Calamante, & Connelly, 2015), COMMIT (Daducci, Dal Palù, Lemkaddem, & Thiran, 2014) or LiFE (Pestilli, Yeatman, Rokem, Kay, & Wandell, 2014) in order to mitigate the limited reliability of streamline count as a proxy of axonal connectivity (Jbabdi & Johansen-Berg, 2011). Given that tractography filtering techniques have non-negligible effects on the topology of structural connectomes (Frigo et al., 2020), an independent analysis is due in order to assess how their use affects the alignability of connectomes.

Notice that in our analysis we used the defined similarity metrics to assess which atlas yields connectomes with higher or lower robustness *in a certain resolution range*. This means that we could not have used the similarity argument to claim that, for instance, the Desikan atlas (68 parcels) should in general be preferred to the Gallardo 1000 atlas. In this sense, we highlight how the considered similarity metrics (GJI, Jr, NMr and Fro) should not be used for selecting the appropriate resolution at which structural connectivity studies should be designed, but they provide a well grounded tool for assessing which of the available atlases at the wanted resolution is most suitable for the considered type of study.

As highlighted throughout the paper, this work analyzes the problems of parcellation selection and brain alignment in the context of *structural* connectivity. Any conclusion we made should not be straightforwardly generalized to functional connectivity or effective connectivity studies, which would require a separate analysis which was out of the scope of this work.

## OPEN SCIENCE

The data and code used in this work are all available at open repositories, as indicated in the text. We uploaded the used code and the obtained connectomes and alignments on the Open Science Framework. They can be found at this link: https://osf.io/depux/.

## ACKNOWLEDGMENTS

The authors would like to thank Dr. Guillermo Gallardo for the help in computing the Gallardo parcellation and Professor Joshua T. Vogelstein for the discussion on the use of FAQ. Also, we are grateful to the OPAL infrastructure from Université Côte d’Azur and Inria Sophia Antipolis - Méditerranée “NEF” computation platform for providing resources and support. This work was funded by the European Research Council (ERC) under the European Union’s Horizon 2020 research and innovation program (ERC Advanced Grant agreement No 694665: CoBCoM - Computational Brain Connectivity Mapping). Data were provided in part by the Human Connectome Project, WU-Minn Consortium (Principal Investigators: David Van Essen and Kamil Ugurbil; 1U54MH091657) funded by the 16 NIH Institutes and Centers that support the NIH Blueprint for Neuroscience Research; and by the McDonnell Center for Systems Neuroscience at Washington University.

## TECHNICAL TERMS

**Parcellation**: subdivision of the brain into distinct regions with respect to morphological, cytoarchitectonic, anatomical, topological or functional criteria.

**Tractography**: method for tracking the trajectory of the axonal pathways that exploits the anisotropy of the diffusion MRI signal.

**Network alignment**: function that maps nodes of a graph onto nodes of another graph, while usually trying to preserve adjacency (endpoints of edges in a graph should map onto endpoints of edges of the other graph).

**Graph similarity**: measure of how much two networks are close with respect to some criteria of topological nature.

**Signature**: feature vector assigned to each node of a graph.

**Bipartite graph**: network whose nodes can be divided in two distinct and non-overlapping sets, such that there are no edges connecting nodes in the same set.

**Bi-stochastic matrix**: matrix of positive entries where each column and row sums to one.

**Breadth-first search (BFS)**: graph traversal that, starting from a root node, explores all its neighbors before moving to the neighbors’ neighbors, and so on.

**Connectome**: the network which encodes the connections of the human brain; it refers to a comprehensive description of the brain’s structural and/or functional connections.

**Structural Connectome**: network that describes the structure of the white matter connections in the brain; it can be obtained via diffusion MRI-based tractography.

**Functional Connectome**: network-like description of the coherence between the activity in different brain regions; it can be obtained by studying co-activation patterns in functional MRI, EEG or MEG.

## Notes

https://osf.io/depux/

### Competing Interest Statement

The authors have declared no competing interest.

### Summary of Updates

The updated version is the one accepted for publication at Network Neuroscience.

