## Supplementary material for "Network alignment and similarity reveal atlas-based topological differences in structural connectomes"

### Supplementary materials for the article
#### *Network alignment and similarity reveal atlas-based topological differences in structural connectomes*

Matteo Frigo[1,*], Emilio Cruciani[2,*], David Coudert[2], Rachid Deriche[1], Emanuele Natale[2], and Samuel Deslauriers-Gauthier[1]

[1]Université Côte d'Azur, Inria, France
{matteo.frigo,rachid.deriche,samuel.deslauriers-gauthier}@inria.fr
[2]Université Côte d'Azur, Inria, CNRS, I3S, France
{emilio.cruciani,david.coudert,emanuele.natale}@inria.fr
[*]These authors contributed equally to this work.

#### A Intra-cohort variability

For each atlas, in Figure 1 we report the boxplot of the distribution of the similarity between the connectomes of each pair of subjects in the studied cohort. The similarity is measured with the proposed Graph Jaccard Index, the Frobenius Norm and the Correlation (cosine distance).

#### B Comparison between WL-align and FAQ

For each atlas and similarity metric, we compared the results obtained with WL-align and FAQ on the 100 subjects in the studied cohort. The statistical significance of these differences is tested with the Wilcoxon signed-rank test at $\alpha = 0.05$. As shown by the figures hereafter presented, all tested differences are statistically significant.

| Metric | Figure |
|---|---|
| Node Matching Ratio | Figure 2 |
| Graph Jaccard Index | Figure 3 |
| J ratio | Figure 4 |
| Frobenius Norm | Figure 5 |

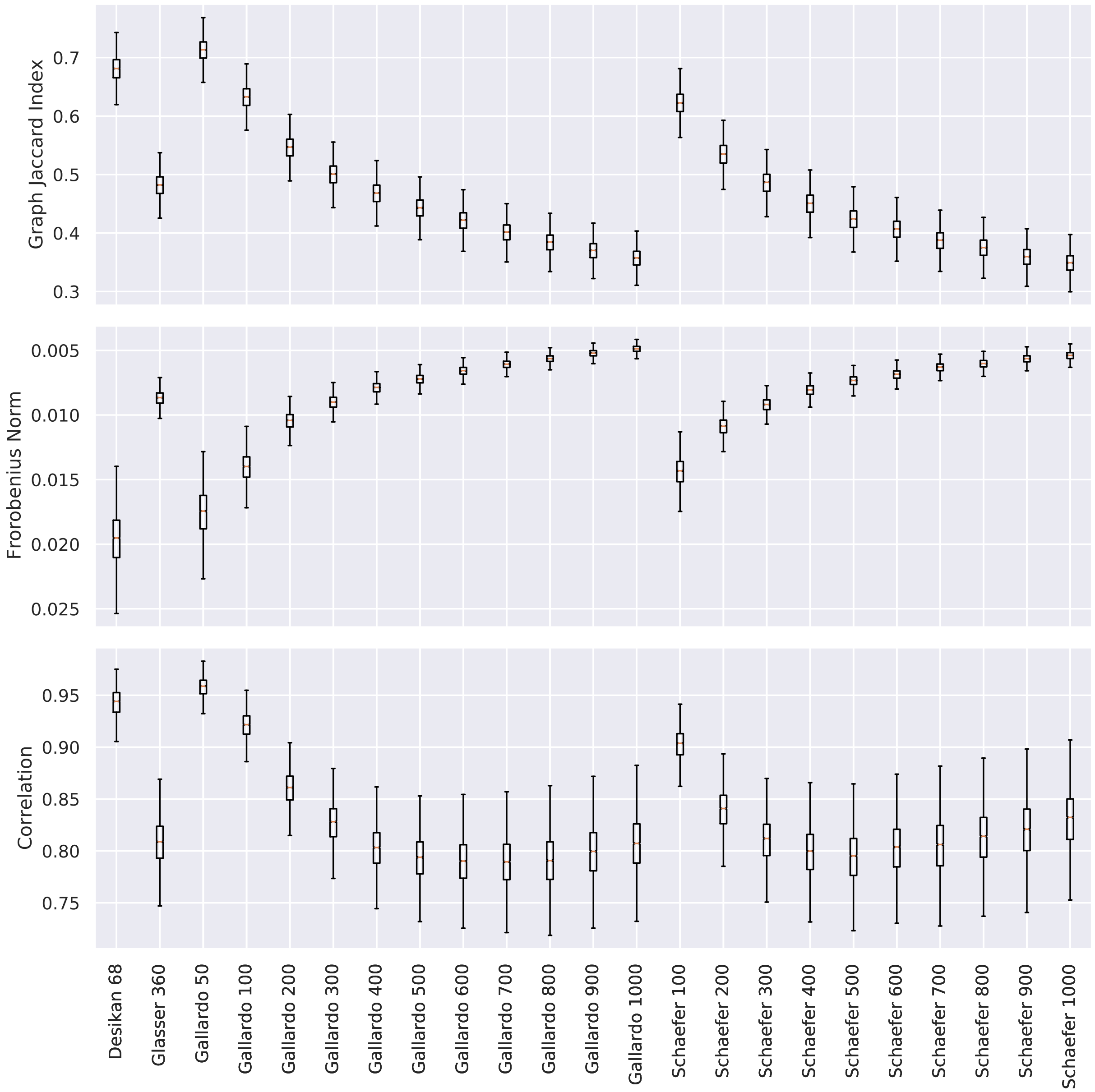


Figure 1: For each atlas, the boxplot of the distribution of the similarity between the connectomes of each pair of subjects in the studied cohort is reported.

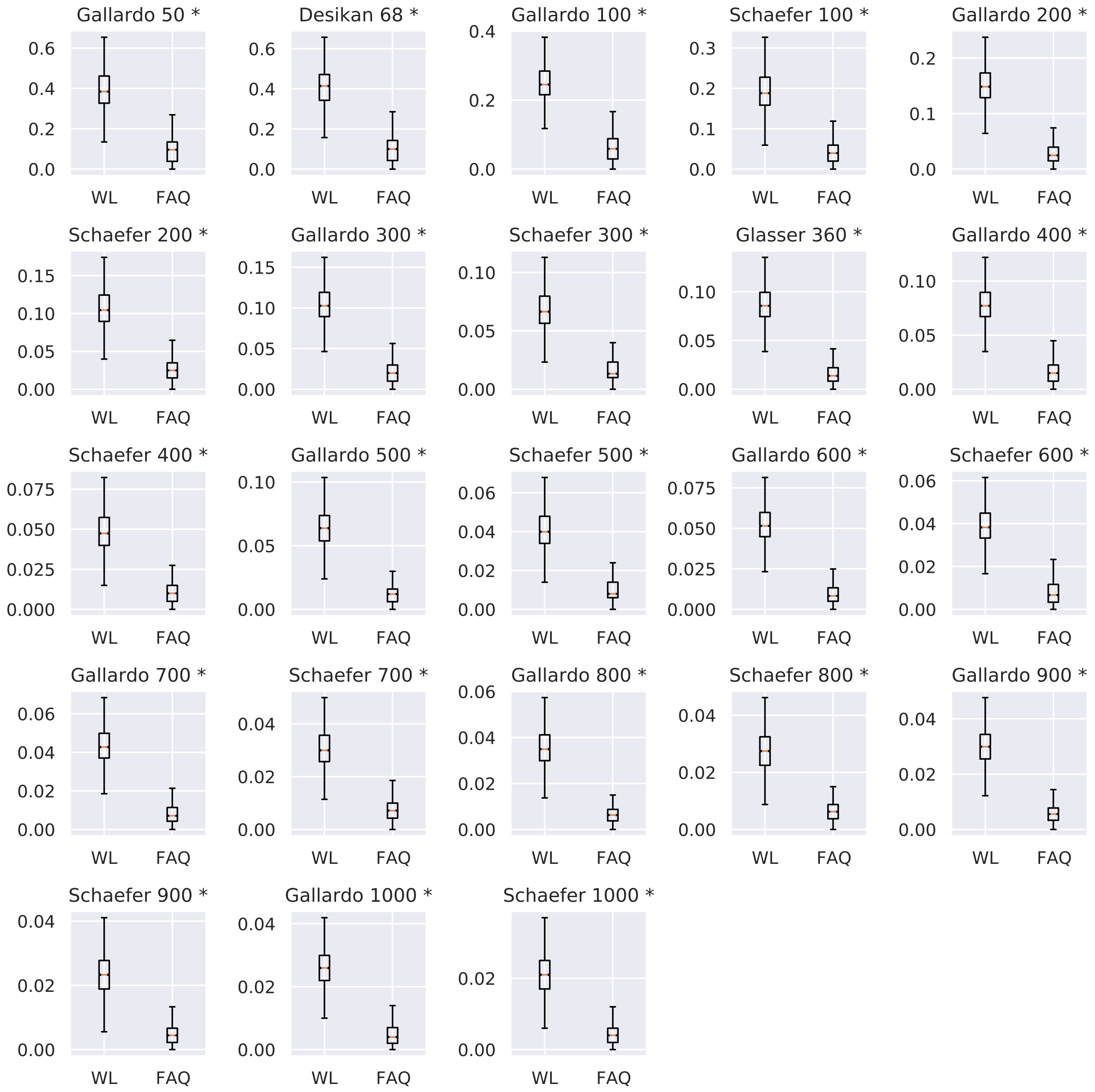


Figure 2: For each atlas, the boxplot of the distribution of the values of the *Node Matching Ratio* obtained with WL-align and FAQ is reported. Differences that are statistically significant ($\alpha = 0.05$) under the Wilcoxon signed-rank test have a * symbol next to the name of the atlas.

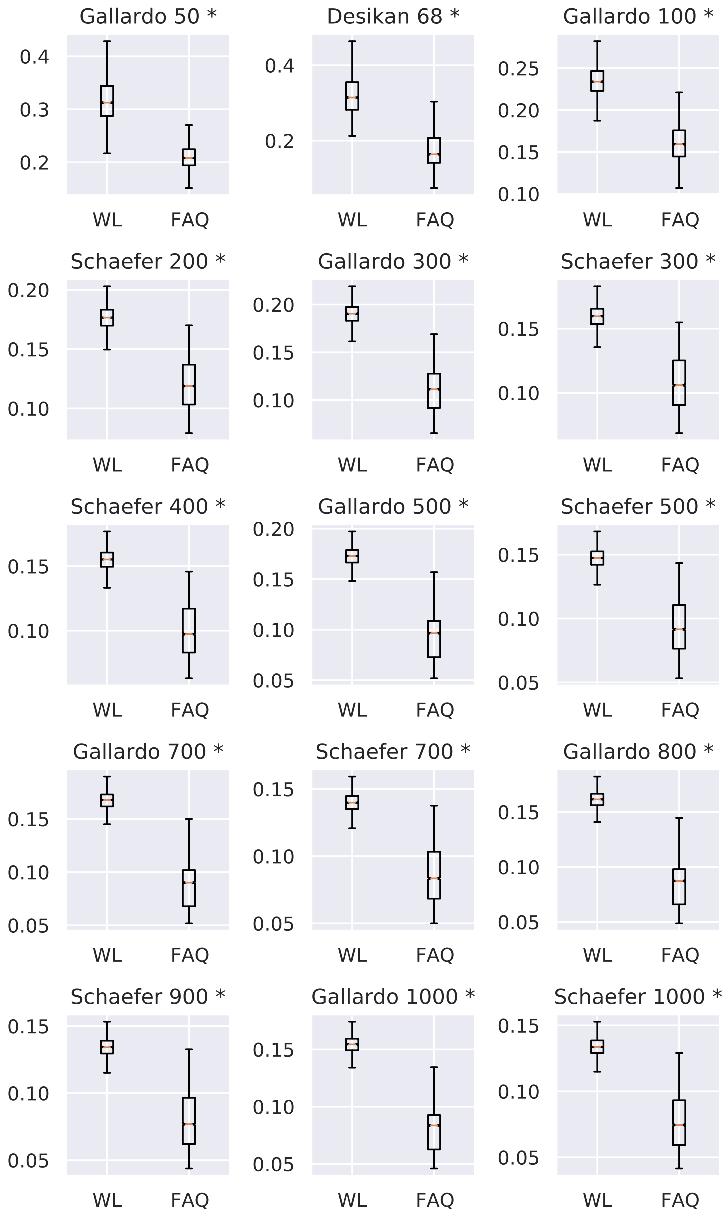


Figure 3: For each atlas, the boxplot of the distribution of the values of the *Graph Jaccard Index* obtained with WL-align and FAQ is reported. Differences that are statistically significant ($\alpha = 0.05$) under the Wilcoxon signed-rank test have a * symbol next to the name of the atlas.

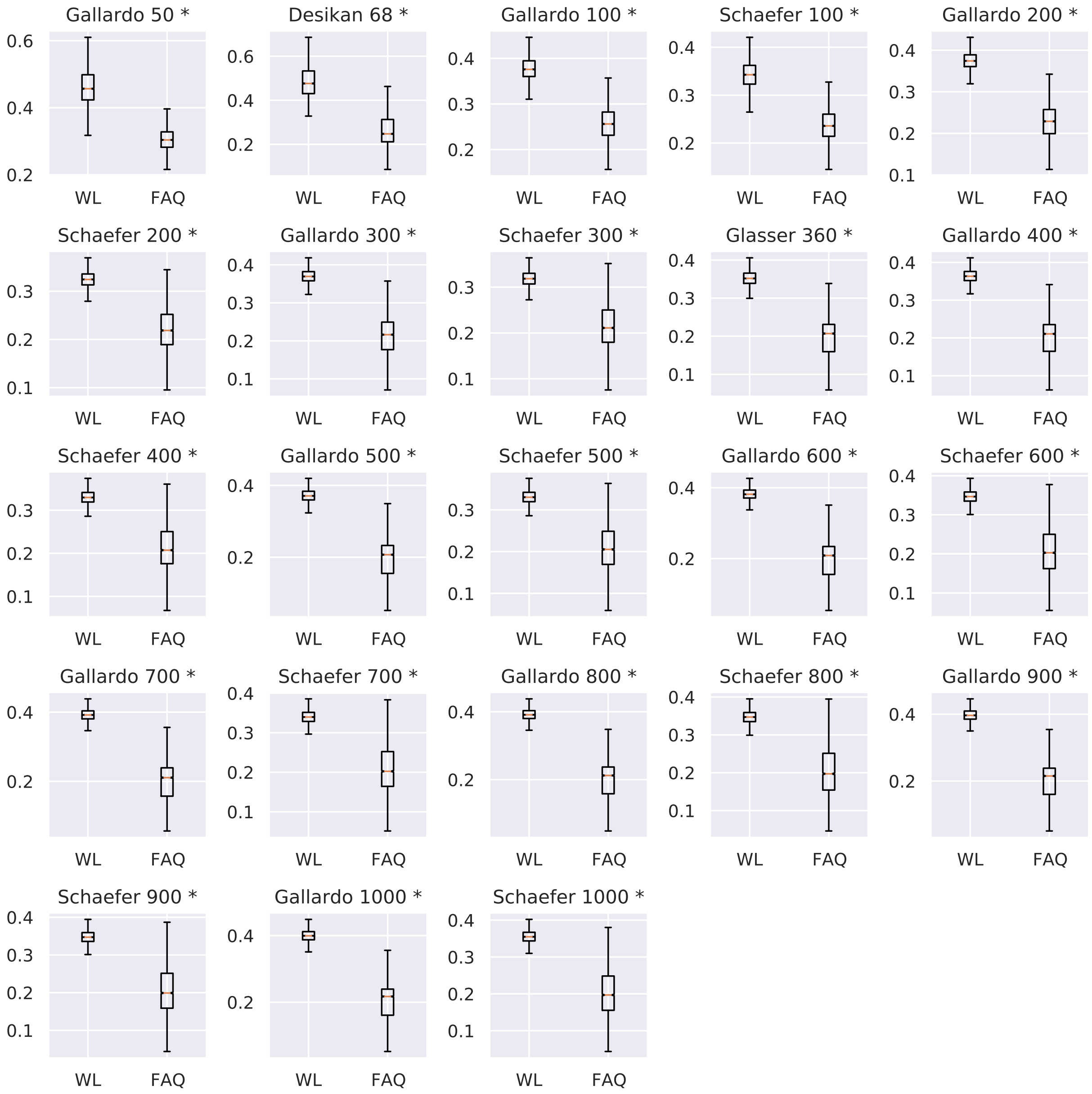


Figure 4: For each atlas, the boxplot of the distribution of the values of the *J ratio* obtained with WL-align and FAQ is reported. Differences that are statistically significant ($\alpha = 0.05$) under the Wilcoxon signed-rank test have a * symbol next to the name of the atlas.

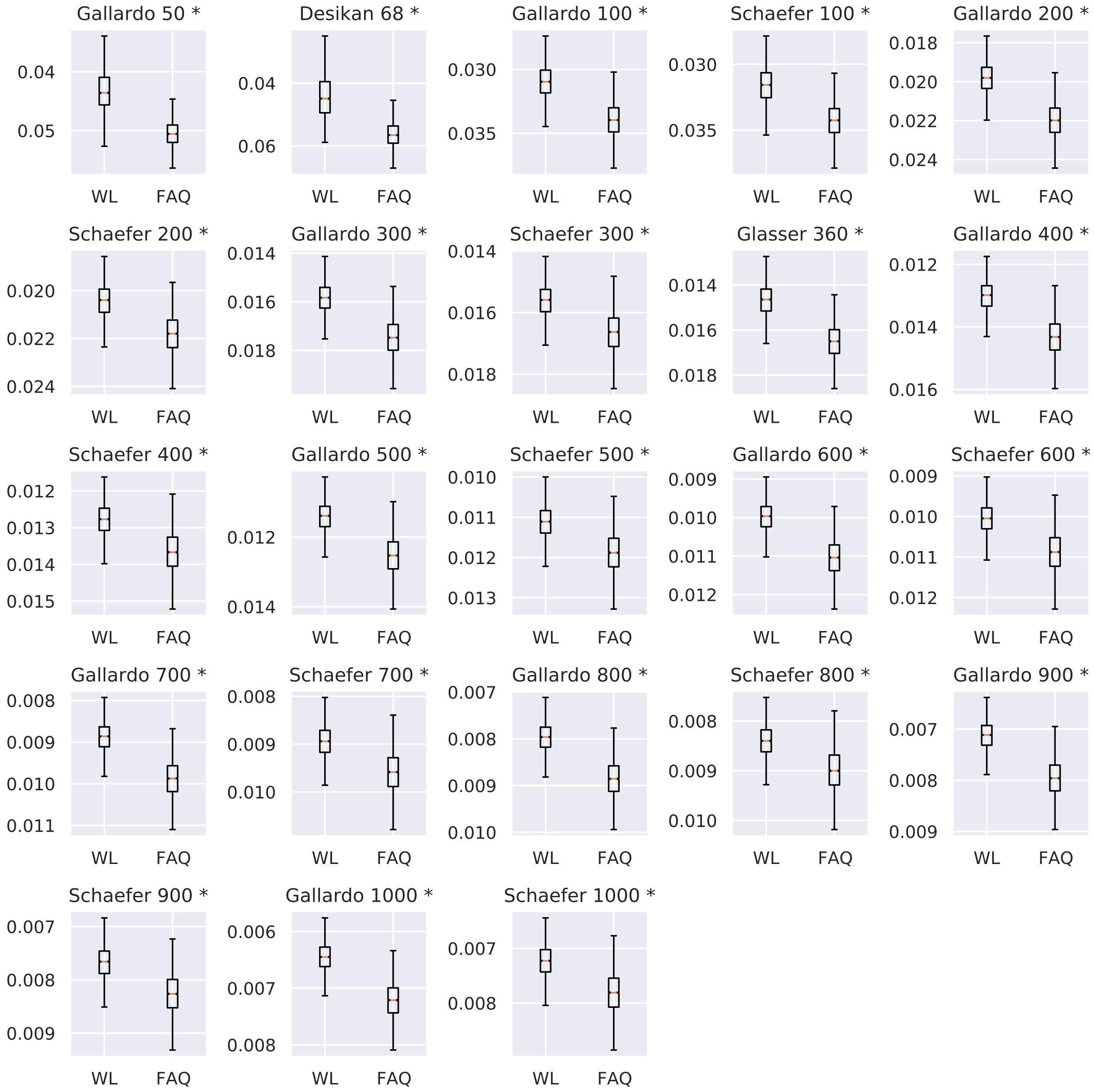


Figure 5: For each atlas, the boxplot of the distribution of the values of the *Frobenius norm* obtained with WL-align and FAQ is reported. Differences that are statistically significant ($\alpha = 0.05$) under the Wilcoxon signed-rank test have a * symbol next to the name of the atlas. The $y$ axis is inverted to reflect the idea that *higher is better*.
